# Fast-evolving alignment sites are highly informative for reconstructions of deep Tree of Life phylogenies

**DOI:** 10.1101/835504

**Authors:** L. Thibério Rangel, Gregory P. Fournier

## Abstract

The trimming of fast-evolving sites, often known as “slow-fast” analysis, is broadly used in microbial phylogenetic reconstruction under assumption that fast-evolving sites do not retain accurate phylogenetic signal due to substitution saturation. Therefore, removing sites that have experienced multiple substitutions would improve the signal-to-noise ratio in phylogenetic analyses, with the remaining slower-evolving sites preserving a more reliable record of evolutionary relationships. Here we show that, contrary to this assumption, even the fastest evolving sites, present in conserved proteins often used in Tree of Life studies, contain reliable and valuable phylogenetic information, and that the trimming of such sites can negatively impact the accuracy of phylogenetic reconstruction. Simulated alignments modeled after ribosomal protein datasets used in Tree of Life studies consistently show that slow-evolving sites are less likely to recover true bipartitions than even the fastest-evolving sites. Furthermore, site specific substitution-rates are positively correlated with the frequency of accurately recovered short-branched bipartitions, as slowly evolving sites are less likely to have experienced substitutions along these intervals. Using published Tree of Life sequence alignment datasets, we additionally show that both slow-and fast-evolving sites contain similarly inconsistent phylogenetic signals, and that, for fast-evolving sites, this inconsistency can be attributed to poor alignment quality. Furthermore, trimming fast sites, slow sites, or both is shown to have substantial impact on phylogenetic reconstruction across multiple evolutionary models. This is perhaps most evident in the resulting placements of Eukarya and Asgardarchaeota groups, which are especially sensitive to the implementation of different trimming schemes.

**Significance Statement:** It is common practice among comprehensive microbial phylogenetic studies to trim fast-evolving sites from the source alignment in the expectation to increase the signal to noise ratio. Here we show that despite fast-evolving sites being more sensitive to parameter misspecifications than mid-rate evolving sites, such sensitivity is comparable, if not smaller, than what we observe among slow-evolving sites. Through the use of both empirical and simulated datasets we also show that, besides the lack of evidences regarding the noisy nature of fast-evolving sites, such sites are of core importance for the reliable the reconstruction of short-branched bipartitions. Such points are exemplified by the variations in the Eukarya+Archaea Tree of Life when subjective alignment trimming strategies are employed.

## Introduction

The suitability of alignment sites for use in phylogenetic inference is influenced by many factors, such as compositional bias, site specific substitution frequencies, and evolutionary rates (1). Tree reconstruction methods often attempt to take these processes into account, either by using more complex evolutionary models, or by removing sites found to violate the assumptions of a simpler evolutionary model, or that are predicted to add phylogenetic noise. One such technique is “slow-fast” analysis, whereby the fastest evolving sites within an alignment are removed to improve the signal-to-noise ratio within the remaining sequence data (2). In the most extreme cases, all but the slowest evolving sites may be removed, and the resulting phylogenies are compared (e.g., Raymann et al. 2015).

The underlying assumption that the slowest-evolving, most highly conserved sites retain the most accurate phylogenetic signal is largely based on concerns about site saturation—that is, multiple substitutions that overwrite earlier substitutions that would otherwise retain phylogenetic information about deep divergences (4). Typical saturation assessment methods are based in comparisons between corrected versus non-corrected pairwise distances among sampled taxa (1, 4, 5). The problematic nature of fast-evolving sites was initially formulated during a time when Maximum Parsimony was the major framework of phylogenetic reconstruction (2, 4, 6), and computing resources constituted a severe bottleneck for modelling its evolution. Even considering that substitution saturation would hamper resolution of phylogenetic relationships between extant taxa, which we discuss as not applicable, such sites may nevertheless contain informative substitutions for resolving relationships between internal nodes of the phylogeny.

The likelihood of a site experiencing a substitution along a branch is dependent upon the bipartition branch length and the rate of substitution inferred for that site (7). Therefore, all things being equal, phylogenetic information regarding short branched bipartitions is less likely to be present among slowly evolving sites. Very slow-evolving sites may not be expected to have experienced any substitutions at all across branches below a certain length threshold, even for a large number of sampled sites. Preferential use of slower-evolving sites may be especially problematic if bipartitions of key interest within a phylogeny have short branches. In such cases, slow-evolving sites may be less likely to retain an informative and accurate signal for resolving these bipartitions when compared to faster-evolving sites. This reasoning calls into question the underlying assumption of “slow-fast” analysis, that, in general, slower evolving sites are more reliable for deeper phylogenetic reconstruction.

Deep Tree of Life phylogenies representing one or more Domains of life are often produced from subsets of highly conserved core protein families, such as riboproteins (3, 8, 9). Is slow-fast analysis appropriate for these datasets? We specifically test the reliability of slow-evolving sites using an aligned, concatenated ribosomal protein dataset, as well as simulated sequence data generated from similar trees and evolutionary models. We find that not only are the slowest evolving sites unreliable and inconsistent in recovering deep phylogenetic signal, but, surprisingly, they are actually less reliable than the fastest-evolving sites, even in the presence of substitution saturation. We specifically show that [1] slow-evolving sites from actual multigene sequence alignments produce trees significantly more phylogenetically incongruent than those produced using fast-evolving sites; [2] slow-evolving sites from simulated sequence data are significantly less likely to recover bipartitions present in the true underlying phylogeny than faster-evolving sites; [3] these effects are strongest among short branched bipartitions; and (4) sites evolving at moderate relative rates consistently out-perform both slow and fast-evolving sites across all metrics. We also show that as taxonomic sampling within a tree increases, internal branches become progressively shorter, making the phylogenetic signal within slow-evolving sites less reliable. From this, we infer that phylogenetic inference from subsampled alignments enriched in slow-evolving sites will become increasingly problematic as sequenced microbial diversity continues to expand.

From these tests, we conclude that removing both the slowest and fastest evolving sites from conserved protein alignments should result in improved phylogenetic resolution for the deepest splits in the Tree of Life, specifically those with short branches. Applying this approach to both real and simulated datasets improves the resolution recovered for many deep bipartitions, improving support for specific evolutionary hypotheses.

## Results & Discussion

### Slow evolving sites contain inconsistent phylogenetic signals in conserved protein datasets

Sites evolving with different substitution rates are expected to display varying degrees of phylogenetic information, as evidenced by consistency of phylogenetic signal. To test this hypothesis, sites from the ribosomal protein alignment published in Hug et al. (2016) were binned into twelve gamma distributed site-specific substitution rate categories (10), and sites from each bin were used to independently reconstruct the tree. Each of the twelve substitution rate categories (SRCs) group sites with similar substitution rates as evidenced by the most likely substitution rate category identified for the site, ordered from the slowest to the fastest-evolving. Invariant sites were omitted before site rate categories were assigned. Sites from each SRC were used to generate Rate Specific Alignment Partitions (RSAPs), from RSAP1 (containing the slowest-evolving sites) to RSAP12 (containing the fastest-evolving sites). Each RSAP was used to generate 1,000 UltraFast Bootstrap (UFBoot) samples (11) using IQTree (12).

The observed phylogenetic signal differs substantially across RSAPs, as demonstrated by the distinct tree-space areas explored by each UFBoot sample (Figure 1). Surprisingly, in addition to being the most dissimilar to the tree generated from the full alignment, UFBoot samples obtained from RSAP1 also display the most inconsistent topologies among UFBoot replicates (Figure S1) as measured by Robinson-Foulds (RF) distances (13). The lesser consistency among topologies obtained from slow-evolving sites, when compared to faster-evolving ones, is likely due to low substitution frequencies not generating enough substitutions along branches to resolve many bipartitions. In fact, the similarity between topologies obtained from each RSAP to the whole alignment phylogeny increases with substitution rates up to RSAP9 (average 1.21 substitutions/site) followed by a subsequent decrease towards the fastest-evolving RSAPs. The inversion of the positive association between substitution rate and similarity to the topology of the whole alignment after RSAP9 suggests an optimal balance between accumulated phylogenetic signal and substitution saturation (**Error! Reference source not found.** and Figure S2a). Importantly, a higher consistency of phylogenetic signal observed for a given RSAP does not necessarily indicate greater phylogenetic accuracy, as the true underlying phylogeny remains unknown. However, phylogenetic consistency is a precondition for construction of a well-supported phylogeny, and so serves as a reasonable metric in the absence of a known true tree, even if such a metric does not account for other possible sources of bias.

**Figure 1.**
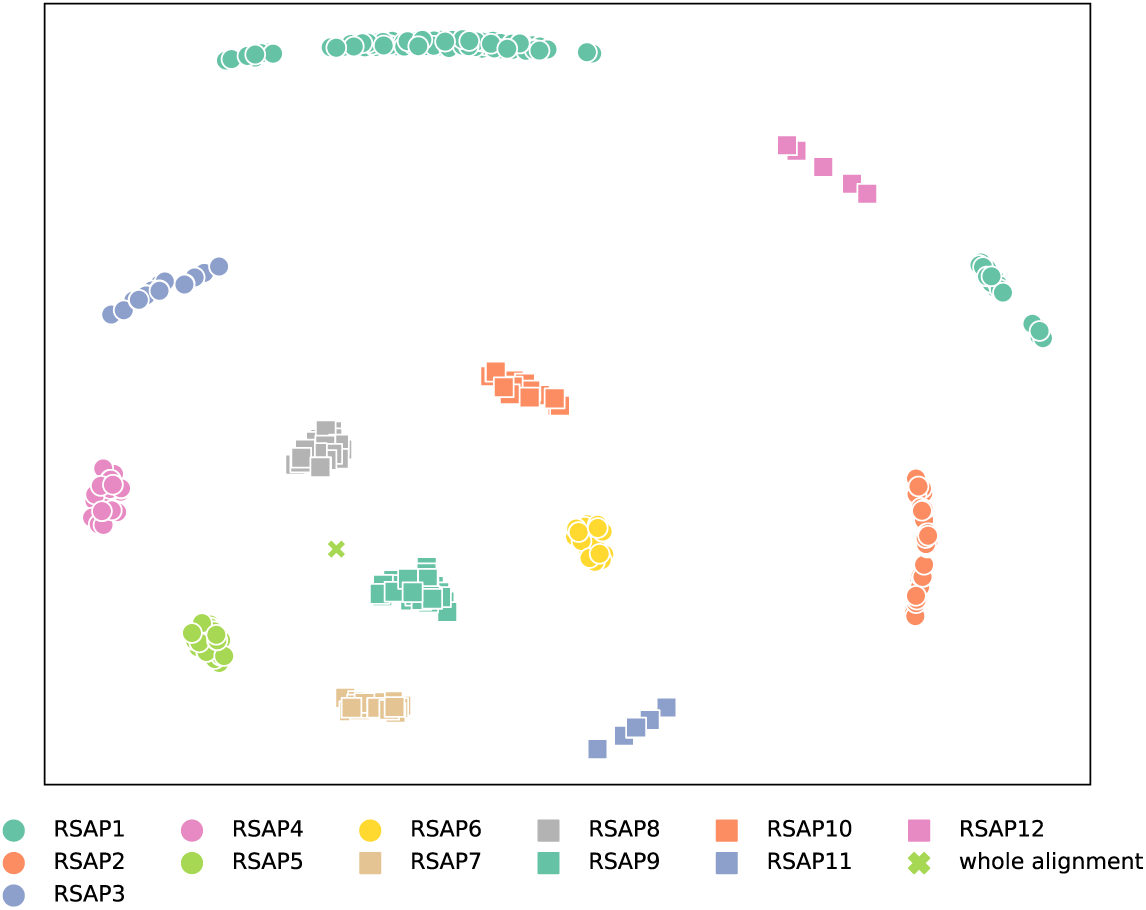
Multidimensional Scalling of RF distances between UFBoot samples and reference topologies. The relative distances between tree topologies within each RSAP informs how sampled trees obtained from a RSAP are consistently similar (e.g., RSAP8 and RSAP9) or dissimilar (RSAP1 and RSAP12) to the overall topology. One hundred trees were randomly chosen from each UFBoot sample. The RF distances between each set of trees, as well as the phylogeny published by Hug et al. was then calculated.

### Short branched bipartitions are less consistently recovered from slow evolving sites

Slow-evolving sites are inherently less likely to experience substitutions along short branches than faster-evolving sites; therefore one would expect slow-evolving sites to less reliably reconstruct short-branched bipartitions. For example, more than half of bipartitions depicted in Hug et al.’s archaeal subtree have branch lengths shorter than 0.05 substitutions/site, corresponding to 129.8 substitutions among its 2,596-site alignment. Since SRC1 possesses an average site-specific substitution rate of 2.332e^−2^ (i.e., accumulating substitutions at a rate 2.332e^−2^ times slower than the average) these sites are expected to experience 0.25 substitutions along a 0.05 branch length (0.194% of the total substitutions characterizing the bipartition). Across the same branch length, sites from SRC12 are expected to accumulate 41.36 substitutions (31.86% of the total substitutions characterizing the bipartition) during the same interval, as their average substitution rate is 3.8243 times faster than the average. Conversely, fast-evolving sites are more likely to undergo substitution saturation along a long-branched bipartition.

Comparing the relative compatibility of bipartitions recovered from each RSAP UFBoot sample to bipartitions found in the tree generated from the full alignment (reference bipartitions), slow-evolving sites are indeed less likely to recover short-branched reference bipartitions. The shorter a bipartition’s branch length, the less likely it is to be consistently recovered by slow-evolving sites (Figure 2a); the faster an RSAP’s substitution rate, the more likely it is to recover shorter branched bipartitions. There is a significant negative Spearman correlation between average site-specific substitution rate and median branch length of reference bipartitions compatible with UFBoot samples (*rho* = −0.972 and *p* = 1.28*e*^−7^). We define compatible reference bipartitions as those present in at least 80% of UFBoot sampled obtained from a specific RSAP.

**Figure 2.**
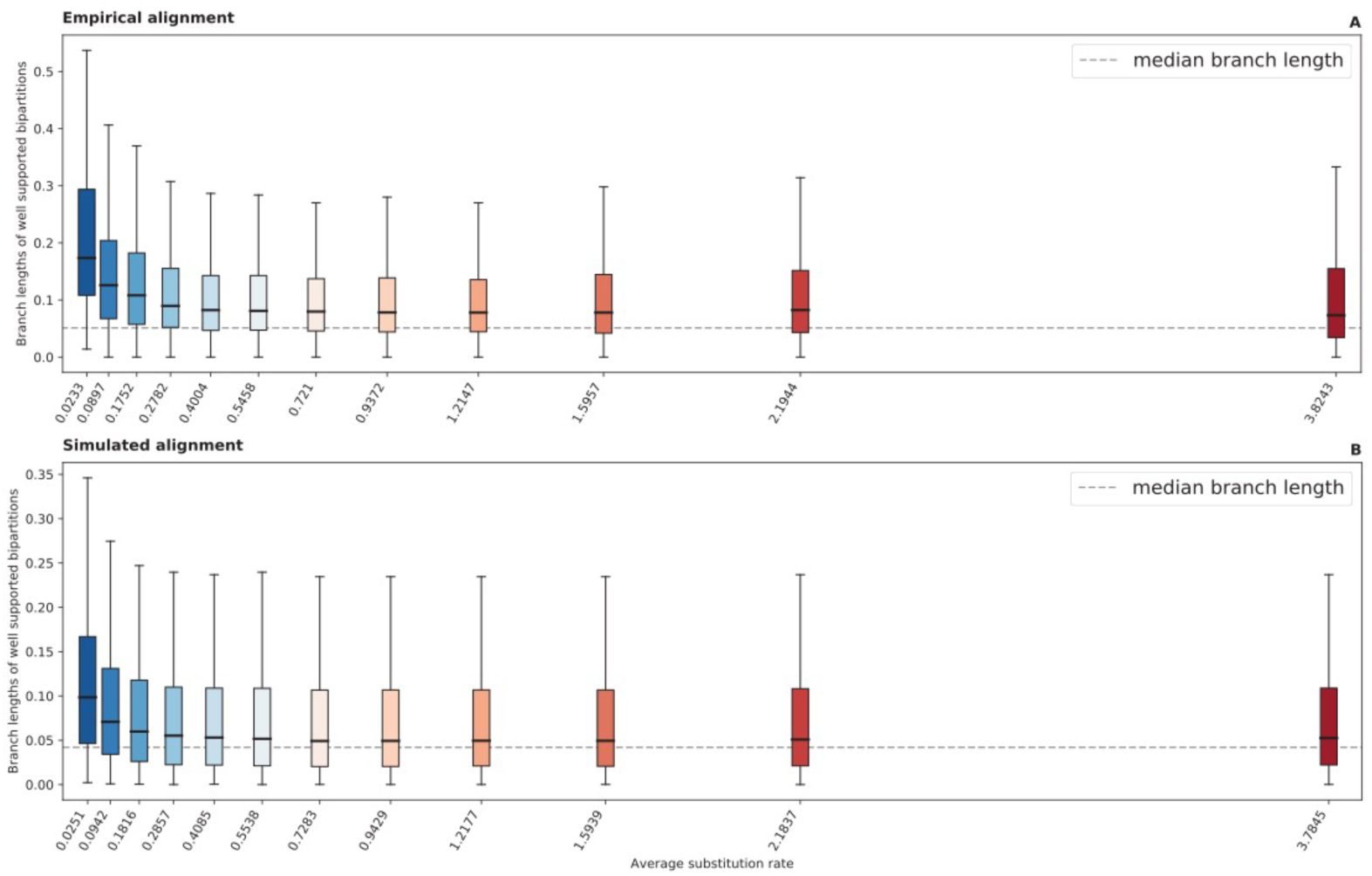
Boxplot distributions of reference bipartitions present in at least 80% of UFBoot samples generated from each RSAP, for both Hug et al. (A) and simulated (B) datasets. The dashed line represents the median internal branch length of each reference phylogeny.

### Slow evolving sites in simulated alignments are less likely to recover true tree bipartitions

Phylogenetic tests of actual sequence datasets can only reliably evaluate consistency across rate categories, rather than accuracy, given that the true underlying tree is inferred rather than known. Therefore, to further assess the accuracy of phylogenetic reconstruction across SRCs, we generated a dataset containing 100 simulated sequence alignments using a known true-tree phylogeny. A random tree topology with 1,000 leaves using a branch length distribution modeled after branch lengths observed within the Hug et al. dataset’s phylogeny was generated and used as a guide tree for sequence simulation (Dataset S1). Simulated alignments were then generated by evolving a random starting sequence using the average amino acid composition and substitution frequencies observed in the Hug et al. dataset. As performed for the Hug et al. dataset’s alignment, sites from each simulated alignment were binned according to twelve gamma-distributed site-specific SRCs, generating twelve RSAPs. Each RSAP from each simulated alignment was then used to generate 1,000 UFBoot tree samples.

The results of the simulated data analyses were in agreement with those observed for the Hug et al. dataset. RF distances between the true tree and UFBoot samples show that slow-evolving sites consistently underperformed in reconstructing the overall true-tree topology when compared with all other faster-evolving RSAPs (Figure 3). Although substantially less pronounced than in RSAPs from the Hug et al. alignment, the deterioration of phylogenetic signal as substitution rates increase is still present among simulated alignments, detected from RSAP10 onward. The slope of a linear regression between average substitution rate and RF distances between the true tree and UFBoot samples represents the pace in which increasing the former impacts the later, positively or negatively. Among the 100 simulated alignments, the average slope of the described regression from SRC9 to SRC12 is 26.23 (Figure S2b), suggesting that although deviation from the true tree still increases in the simulated scenario, it does so 45 times slower than observed among equivalent SRCs from the Hug et al. dataset (Figure S2a).

**Figure 3.**
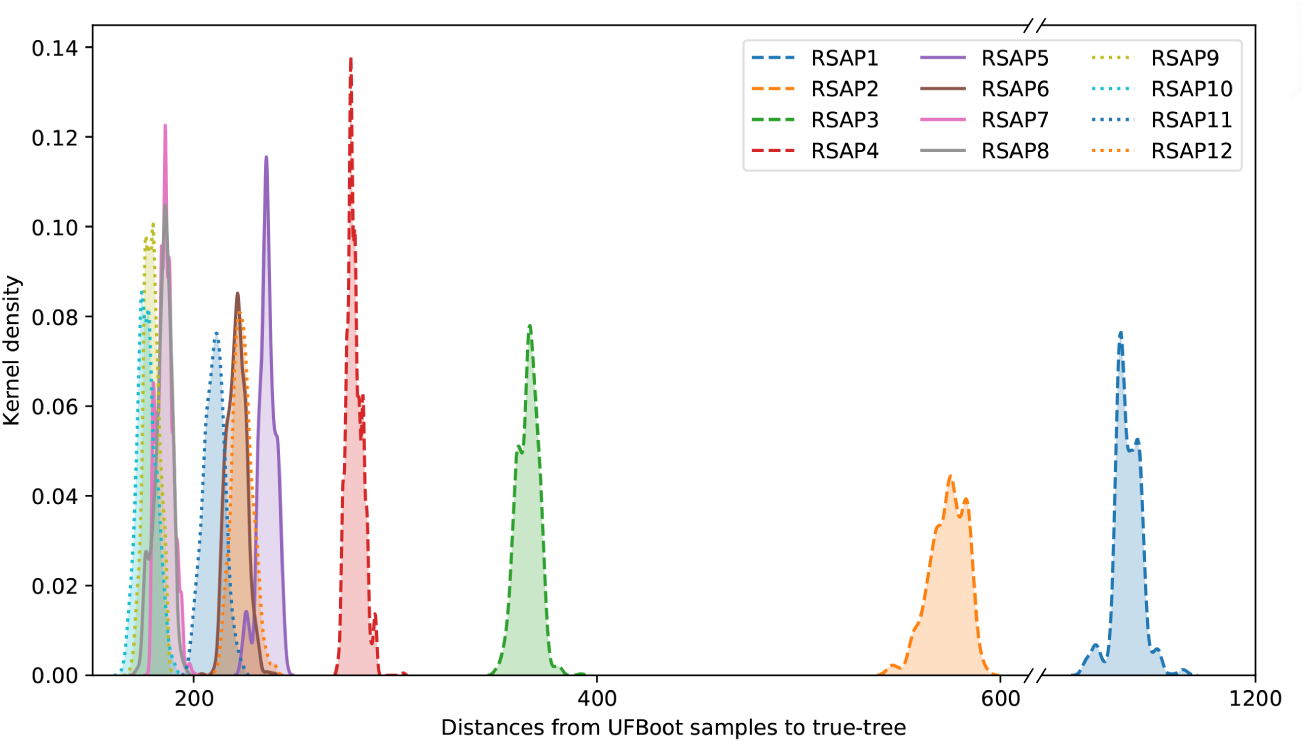
Distribution of RF distances between RSAP UFBoot samples and the true tree topology. Distances plotted are all derived from a single simulated alignment replicate, as distributions are consistent across all 100 replicates.

### Slow evolving sites in simulated sequence datasets are biased against reconstructing true short-branched bipartitions

A further breakdown of true bipartition recovery frequencies from simulated RSAPs shows that these are not independent of site rates. 91 out of the 100 simulated alignments showed significant negative Spearman correlations between average substitution rate and the branch length of bipartitions present in at least 80% of UFBoot samples (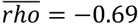 and *q* < 0.05). True tree bipartitions with shorter branch lengths were more frequently reconstructed by faster-evolving than by slower-evolving sites, as shown by the fastest and slowest-evolving RSAPs respectively reconstructing 73% and 18% of bipartitions with branch lengths below the median (Figure 2b). Interestingly, there was no detected correlation between reconstructing true bipartitions with long branches across substitution rate categories. Remarkably, UFBoot samples from all simulated RSAPs were consistent in accurately recovering ~99% of the top 25% longest branched bipartitions. This suggests that [1] given the large absolute number of substitutions present in such bipartitions they can be found even among the slowest-evolving sites; and [2] substitution saturation in fast-evolving sites does not necessarily significantly obscure phylogenetic signal, or impair phylogenetic reconstruction, even along long branches where many such substitutions are expected to occur.

### Substitution saturation does not explain the loss of phylogenetic signal from fast-evolving sites

The minor deterioration of phylogenetic signal among fastest-evolving sites from simulated alignments is unexpected given the high degree of substitution saturation present in simulated alignments (Figure S2b), especially when compared to the much more dramatic deterioration observed for the Hug et al. dataset. Since branch lengths and substitution rate categories from simulated datasets were modeled after the Hug et al. dataset, saturation by itself cannot be the major force driving the deterioration of phylogenetic signal among the fastest-evolving sites in real sequence data. The concept of substitution saturation causing loss of phylogenetic signal has been questioned before using small datasets and limited taxonomic sampling (6). Alternatively, these results suggest that the consistent deterioration of phylogenetic signal among the fastest-evolving sites is more likely to be due factors not replicated in the simulated dataset. These potentially include alignment errors (14) and/or the fitting of a sub-optimal substitution model (15–17). These two possibilities were further investigated.

Regardless of the phylogenetic information contained in an aligned site, fast-evolving sites are more prone to misalignment simply due to an increased number of states shared by a subset of taxa at nonhomologous sites (14). As both ancestral relationships between taxa and their internal sequence states are unknown variables in phylogenetic reconstruction, it is impossible to directly assess alignment accuracy of empirical datasets. However, given the greater diversity of alignment variations possible within gap-rich regions, sites flanked by gap-rich regions should be enriched in misaligned residues. As such, these were used as a proxy for poorly aligned sites. Conversely, sites within blocks of sequence with few gaps are less likely to be misaligned given fewer combinations of similarly scored local alignment variations. By restricting sampling of fast-evolving sites (from SRC10 to SRC12) exclusively to sites flanked by ungapped regions of the alignment, the resulting RSAPs are expected to contain substantially fewer misaligned sites, and therefore more closely match the profile observed for simulated sites evolving at similar rates. Indeed, the generated UFBoot samples from these RSAPs became significantly more similar to the reference tree (Figure 4). Furthermore, as expected, restricting fast-evolving site sampling to sites flanked by gap-rich regions resulted in UFBoot sample phylogenies even more dissimilar to the reference tree. This result supports that misalignment plays a significant role in the loss of phylogenetic accuracy among fast-evolving sites, regardless of saturation effects.

**Figure 4.**
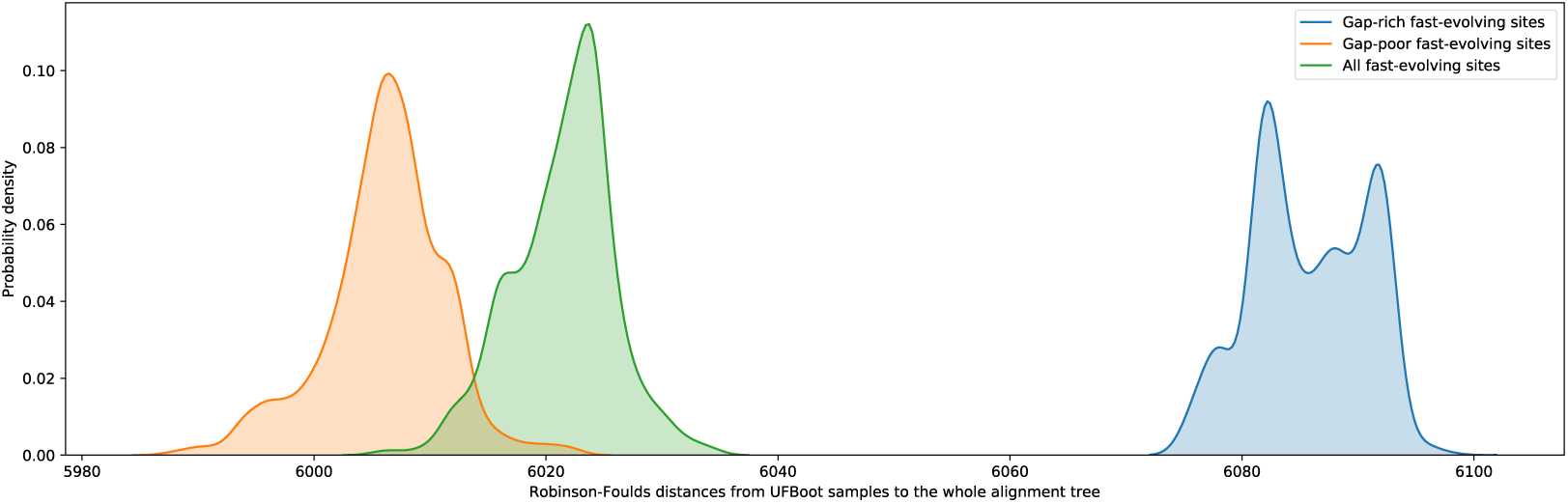
Distribution of Robinson-Foulds distances between the phylogeny proposed by Hug et al. and UFBoot samples obtained using three subsets of fast-evolving sites, SRC10 to SRC12: [1] fast-evolving sites flanked by any number of gaps in green, [2] fast-evolving sites flanked by gap-rich regions in blue, [3] fast-evolving sites flanked by gap-poor regions in orange. Gap-rich regions were defined as containing 2,000 gaps or more, while gap-poor regions as containing 500 or less gaps. 105 sites from distributions 1 and 2 were randomly selected and used to generate UFBoot samples in order to match the number of fast-evolving sites flanked by gap-rich regions.

The impact of sub-optimal substitution models was also assessed. Trees were generated from simulated alignments using a model different than the one the alignments were generated from, specifically, the Dayhoff model (18) instead of LG (19). Sites from simulated alignments were binned into SRCs using the Dayhoff substitution model, followed by RSAP construction, and UFBoot replicates were also generated under the Dayhoff model. UFBoot replicates generated under the Dayhoff model showed a significant increase in the average slope of linear regressions (*q* < 0.05, Figure S3), suggesting that fast-evolving sites are more sensitive to poorly fitted substitution models, contributing to inaccuracy of phylogenetic reconstruction. Interestingly, UFBoot replicates from slow-evolving sites (SRC1 to SRC3) reconstructed under the Dayhoff model were more similar to the true tree than those reconstructed under the “true” LG model (Figure S3). These results might be explained by the cumulative impact of substitution probabilities. The more substitutions a site experiences along its history, the more impactful small changes in the model tend to be, which also increase overfitting related errors, as more events are fitted into a single site (15–17).

Based on these results, we conclude that substitution saturation alone is not the main cause of phylogenetic inaccuracy observed for fast-evolving sites. Rather, it is likely that an increased frequency of indels amongst the fastest-evolving sites lead to misalignment in some sequence regions, and to poor model fitting having a greater impact. Both of these processes may therefore obscure the potential for these sites to reconstruct phylogenetic information despite sequence saturation effects.

### Phylogenetic reconstructions using rate-specific subsets of sequence alignment data

The relatively short internal branch lengths of bipartitions in the phylogeny reported by Hug et al. (e.g., median branch length of 0.05 substitutions/site) is likely related to the general underperformance of slow-evolving sites in recovering this topology. The results from simulated alignments suggest that objective alignment trimming strategies based on site-specific substitution rate categories could improve phylogenetic reconstruction from this dataset, as well as other comparable datasets. In order to test the impact of these strategies, three distinct site trimming approaches were used in reconstructing the phylogeny of the archaeal and eukaryal subtree from the Hug et al. dataset: [1] trimming slow-evolving sites only, from SRC1 to SRC4 (Figures 5b, 5f, and 5j); [2] trimming fast-evolving sites only, from SRC10 to SRC12 (Figures 5c, 5g, and 5k); and [3] trimming both slow and fast-evolving sites (Figures 5d, 5h, and 5l). Alignment partitions resulted from trimming either slow or fast-evolving sites contain 1,886 and 2,130 sites, respectively (72.6% and 82% of the whole alignment). While trimming both slow and fast-evolving sites led to an alignment partition with only the 1,442 sites binned from SRC5 to SRC9, corresponding to 55% of the whole alignment (Dataset S2). Alignments resulting from each trimming strategy, together with the whole alignment, were then used for phylogenetic reconstruction under three distinct substitution models: LG+G, the C60 mixture model, and the distribution free LG4X. Major relationships within the archaeal/eukaryal tree were significantly impacted by these test conditions (Figure 5). For example, the grouping of Asgardarchaeota with Eukarya is recovered in only five of the assessed combinations: the whole alignment under any substitution model (Figures 5a, 5e, and 5i), trimming fast-evolving sites under the LG+G model (Figure 5c), and by trimming both slow and fast-evolving sites under the C60 mixture model (Figure 5h), although not highly supported in any case. The remainder of the reconstructed phylogenies, seven out of twelve, presented Eukarya as sister to the TACK superphylum, and Asgardarchaeota as sister to the Eukarya+TACK clade, a topology which has also been recently proposed using a distinct dataset (20). The only combination of alignment and tree reconstruction model to recover a well-supported (i.e., UFBoot > 95%) placement of Eukarya is found by trimming both slow and fast-evolving sites under the LG+G model. In this case, the TACK superphylum is sister to Eukarya (Figure 5d).

**Figure 5.**
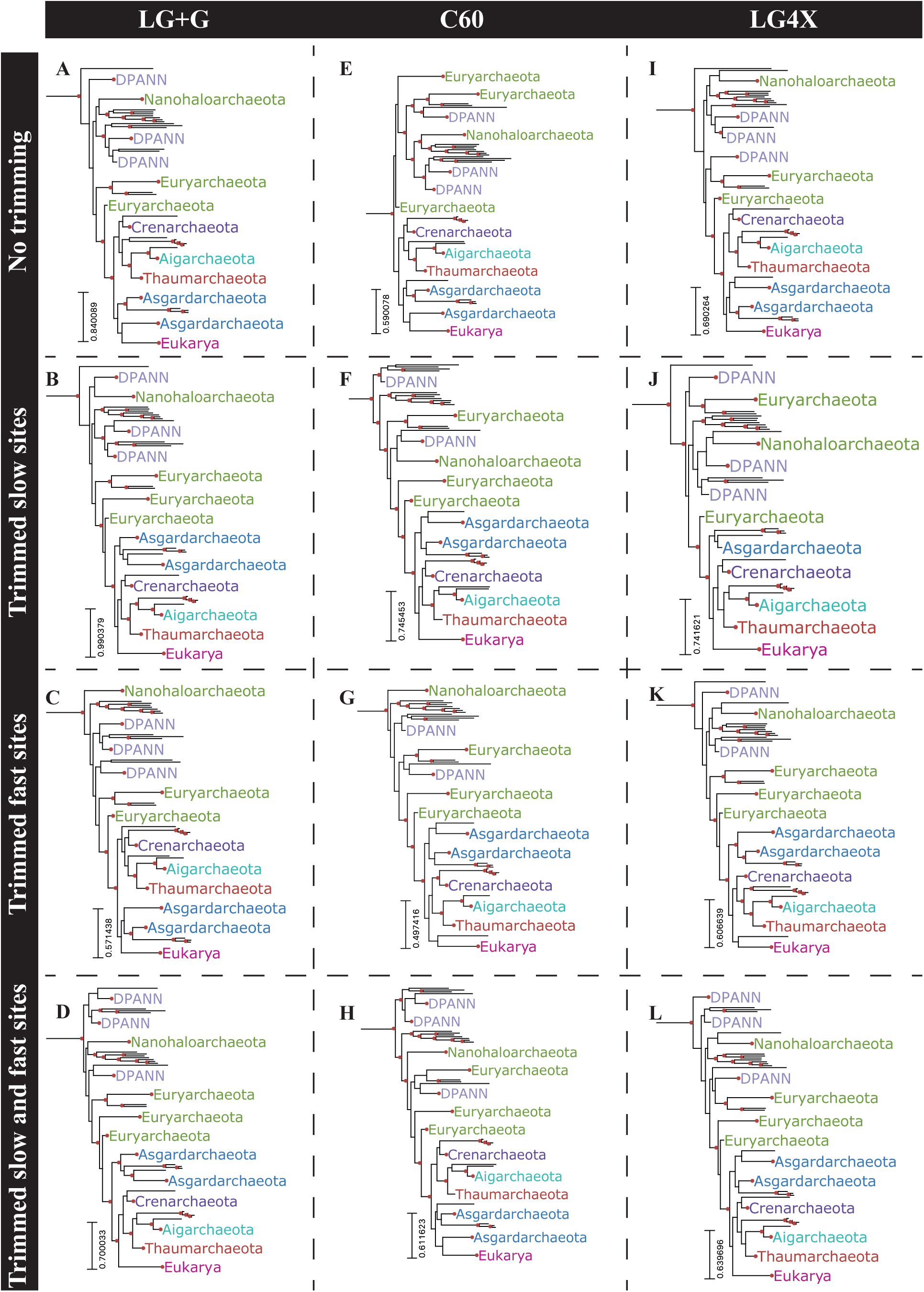
Archaea+Eukaryota subtrees obtained by submitting four distinct alignment partitions of the Hug et al. dataset (whole alignment: A, E, and I; trimmed slow-evolving sites: B, F, and J; trimmed fast-evolving site: C, G, and K; and trimmed both slow and fast-evolving sites: D, H, and L) to three distinct substitution models (LG+G: A, B, C, and D; C60: E, F, G, and H; and LG4X: I, J, K, and L). Highlighted bipartitions have a UFBoot support greater than 95%.

The combination of short internal branch lengths within the Archaea+Eukarya clade with the extremely long branch leading to Bacteria, and the relatively small number of concatenated aligned proteins (16 ribosomal proteins) prevent a realistic expectation of this dataset recovering an accurate rooting for Archaea+Eukarya (20, 21). Nevertheless, recovered rootings to terminal branches remain more suspect, as they are more likely to represent long branch attraction artifacts in the absence of true phylogenetic signal (22). Phylogenies reconstructed from alignment partitions with both slow and fast-evolving sites trimmed led to deeper Archaea+Eukarya roots when reconstructed under LG+G or LG4X substitution models (Figures 5d and 5l). Under the C60 substitution model this alignment partition resulted in a somewhat shallower rooting (Figure 5h) when compared to the whole alignment (Figure 5e), although still recovering a deeper root than from trimming either the faster or slower sites (Figures 5f and 5g).

### Phylogenetic impact of fast-evolving sites within gap-rich regions

In the light of previously discussed results (i.e., substitution saturation not leading to deterioration of phylogenetic signal among fast-evolving sites) we reconstructed the Hug et al. dataset phylogeny by trimming only fast-evolving sites flanked by gap-rich sites from the whole alignment. The 249 sites fulfilling both requirements, fast-evolving and within gap-rich regions, were trimmed from the whole alignment and the resulting alignment partition submitted for reconstruction under three evolutionary models: LG+G, C60, and LG4X (Figure 6). Reconstructions using all three substitution models reported the TACK superphylum as sister to Eukarya, with high bipartition support (93% UFBoot support) under the LG+G model (Figure 6a). Asgardarcheota was reported as sister to the Eukarya+TACK clade in by all three reconstructions.

**Figure 6.**
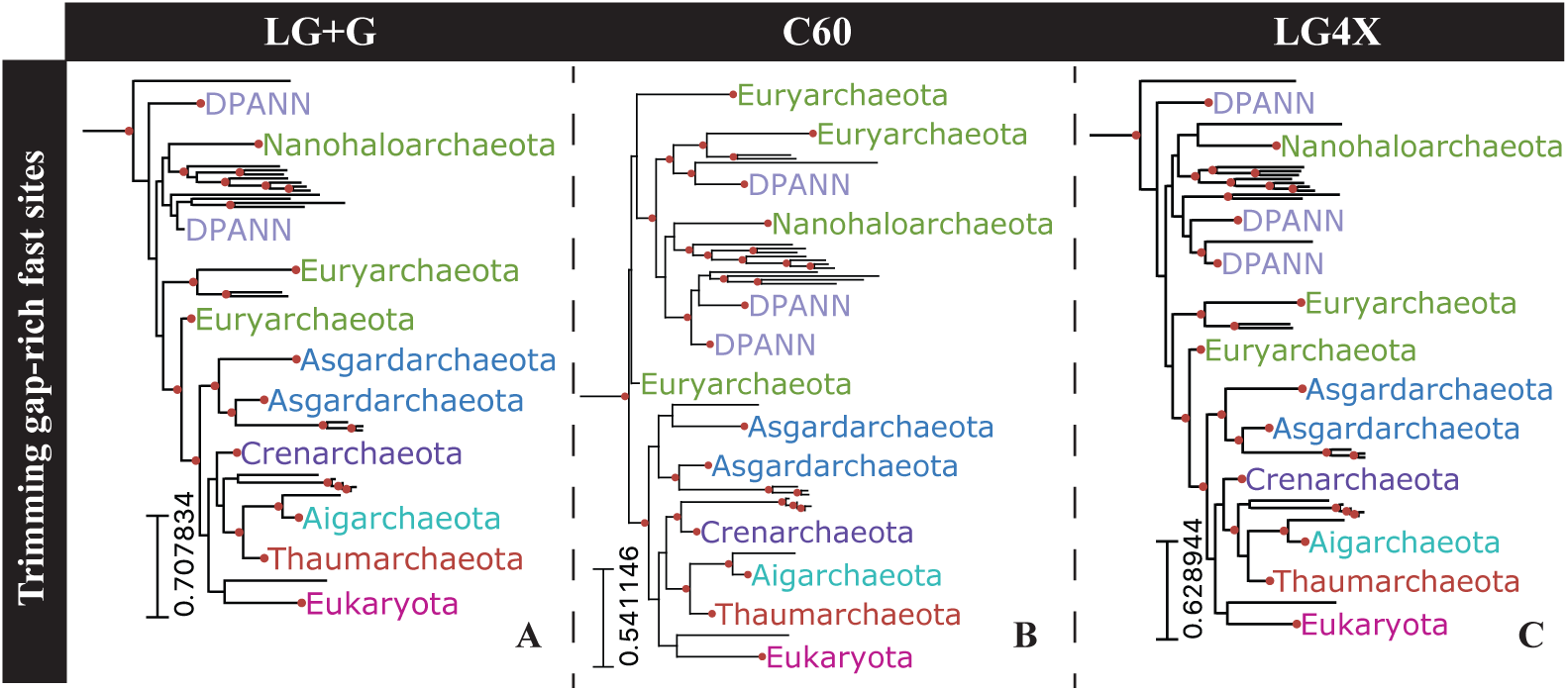
Eukarya+Archaea subtrees reconstructed by removing 249 fast-evolving sites flanked by gap-rich sites from the Hug et al. dataset. Topologies were reconstructed under three distinct substitution models: (A) LG+G, (B) C60, and (C) LG4X. Highlighted bipartitions have a UFBoot support greater than 95%.

### Composition heterogeneity among substitution rate categories

Differences in phylogenetic consistency observed across RSAPs may also be driven by site rate-specific compositional biases, especially when such biases violate assumptions of substitution models. Sites evolving at varying rates may adapt to changes in the underlying mutation bias with varying efficiencies (23). The slowest evolving sites do indeed show substantial compositional bias when compared to the whole alignment (Table 1). Such compositional bias is not uniformly distributed among all twenty amino acids, as highlighted by the massive enrichment of glycine within RSAP1 sites (Figure S4). The distinct biases detected among RSAPs leads to their amino acid frequencies being better modeled by distinct combinations of substitution probabilities (Dayhoff, JTT, WAG, and LG) and amino acid frequencies (default and empirical) when assessed using homogeneous substitution rates (Table 1).

**Table 1.** Best-fitting substitution models of each RSAP from the Hug et al. dataset.

|  |  |  |  |  |  |  |  |  |  |  |  |  |
| --- | --- | --- | --- | --- | --- | --- | --- | --- | --- | --- | --- | --- |
| RSAP | 1 | 2 | 3 | 4 | 5 | 6 | 7 | 8 | 9 | 10 | 11 | 12 |
| Average site-specific substitution rate | 0.023 | 0.089 | 0.175 | 0.278 | 0.4 | 0.545 | 0.721 | 0.937 | 1.214 | 1.595 | 2.194 | 3.824 |
| SSE of aa compositional bias | 0.14 | 0.012 | 0.003 | 0.002 | 0.003 | 0.003 | 0.003 | 0.006 | 0.005 | 0.009 | 0.01 | 0.01 |
| Best-fit substitution model | WAG | LG | LG | LG+F | LG+F | LG+F | LG | LG | LG | WAG | WAG | WAG |

\* in comparison to the whole alignment.

While empirical amino acid frequencies may be a better overall fit to alignment data in some cases, if the bias is largely provided by the least informative sites, adopting empirical frequencies may not be the optimal approach. Applying empirical or default amino acid frequencies from alignments with either slow or fast-evolving sites trimmed produced substantial changes in the reconstructed topology, as measured by RF distances between UFBoot samples (Figure S5). Trimming both slow and fast-evolving sites reconstructed topologies significantly more robust to amino acid frequency changes (Figure S5) despite these alignments containing fewer sites. When jackknifed to the same alignment length as the partition with trimmed slow and fast-evolving sites, eight out of ten jackknives from the whole alignment were shown to be more sensitive to amino acid frequency changes than the alignment partition with trimmed slow and fast-evolving sites (Figure S6). Using default or empirical amino acid frequencies with the full alignment substantially impacts phylogenetic reconstruction for the dataset, respectively recovering either 2-Domain or 3-Domain topologies (Figure 7).

**Figure 7.**
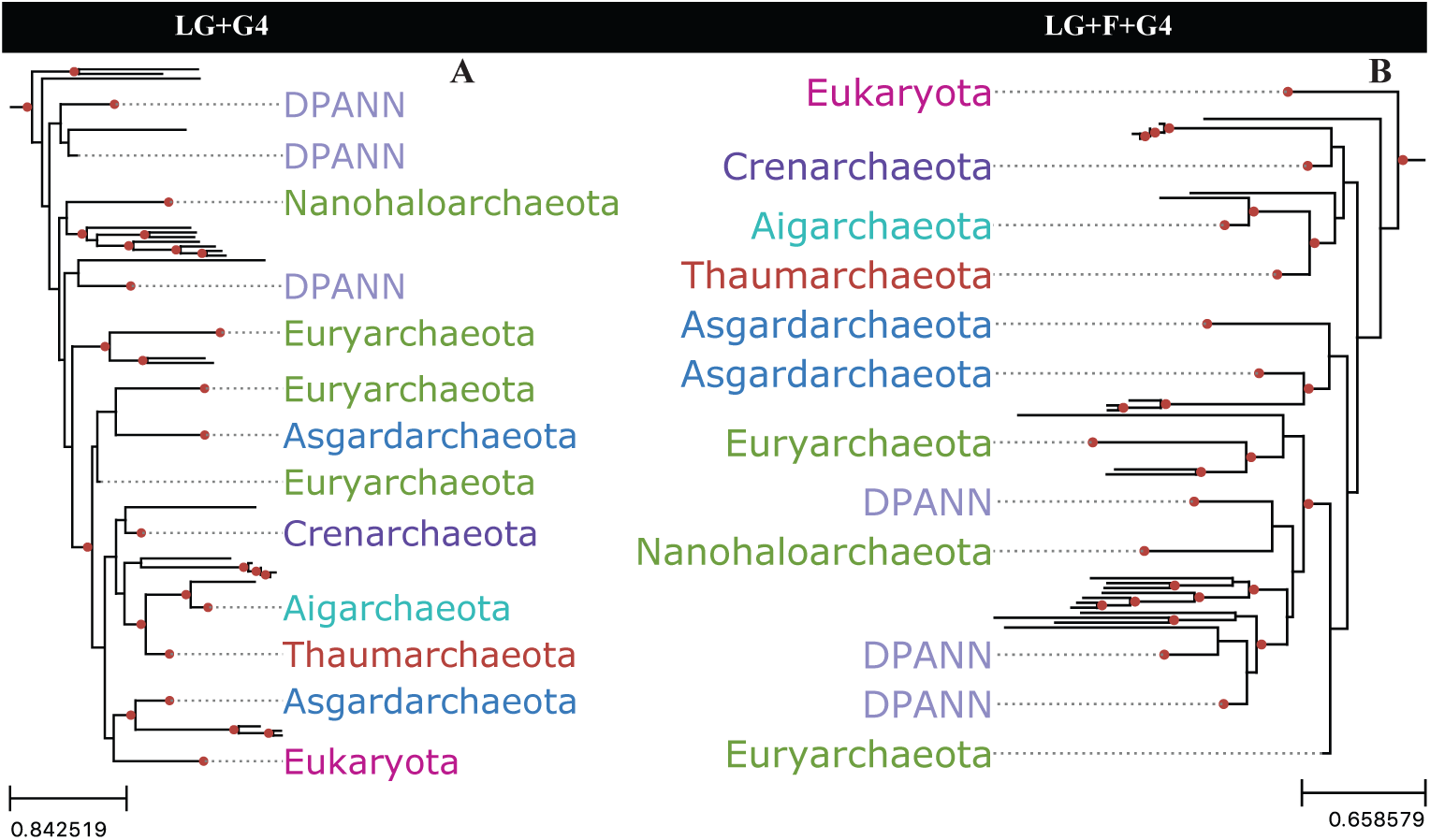
Archaea+Eukarya subtrees reconstructed while varying amino acid frequency in the substitution model. A) Tree reconstructed using equilibrium frequencies default to the LG substitution model. B) empirical state frequencies as estimated from the alignment. Nodes with UFBoot support 95% or greater are highlighted by red circles.

### Short deep branches increase in frequency with increased taxon sampling

Branch lengths are representations of evolutionary distances between two sequence states; in the case of internal bipartitions, this distance is effectively the distance between two inferred ancestors of the sampled taxa. A bipartition with a long branch length is often a consequence of poor sampling within a group (24). This may be caused by patterns of extinction, in which case intermediates do not exist to be sampled; the inability to sample unknown or unsequenced lineages; or deliberate down-sampling of taxa for tractable phylogenetic analysis or taxon sampling balance. Improving taxonomic sampling within a group consequently increases the number of reconstructed intermediary ancestors (nodes) within it. The more intermediary nodes reconstructed between taxa, the more closely related these will be, leading to shorter branch lengths in the reconstructed phylogeny.

Given the performance of fast-evolving sites in recovering short branched bipartitions, the utility of fast-evolving sites in resolving phylogenies should therefore increase as more genomes are sequenced and coverage of genomic diversity becomes denser, as this increases the predominance of short branch bipartitions in phylogenies. To illustrate this, branch lengths were re-estimated using random subsamples of the Hug et al. dataset. A clear downward trend in branch length was observed as taxon sampling increased. Sample sizes were gradually increased from 10% up to 90%, each replicated 100 times (Figure 8). The negative correlation between branch length and sample size emphasizes the importance of better characterizing the impact of rate-specific site partitioning in phylogenetic reconstruction as taxonomic coverage continues to increase.

**Figure 8.**
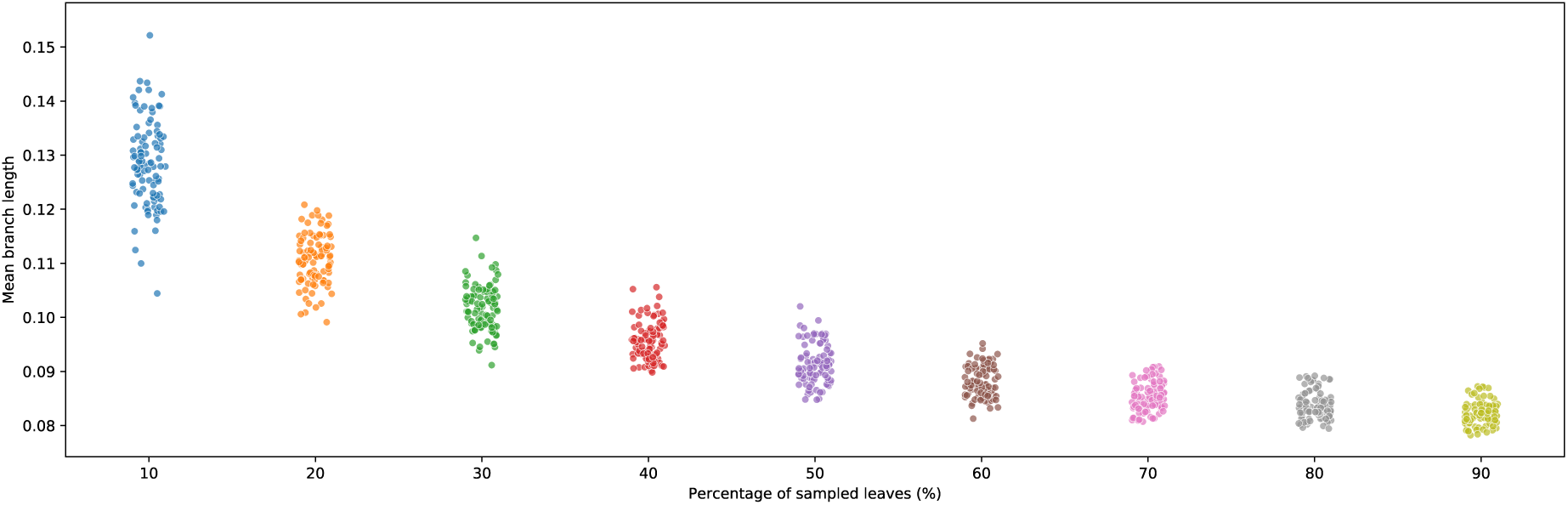
Average re-estimated branch lengths from 100 random taxonomic subsamples of leaves present in Hug et al.’s published phylogeny. Subsampling was increased by 10% at each step.

## Conclusions

It is common practice among microbial phylogenetic studies to trim fast-evolving sites based on substitution saturation metrics obtained using Slow-Fast analysis (3, 25, 26). Testing the assumptions underlying this methodology using both real and simulated datasets based on ribosomal protein alignments and Tree of Life-level sequence diversity, it is clear that, perhaps counterintuitively, fast-evolving sites overperform slow ones with respect to consistency and accuracy of phylogenetic reconstruction. The poorer performance of slow-evolving sites is especially apparent in reconstructing bipartitions with short branch lengths. When compared to sites from the middle of the substitution rate spectrum, fast-evolving sites do show a significant loss of accuracy during phylogenetic reconstruction using both empirical and simulated sequence alignment datasets. However, comparisons using empirical and simulated datasets show that misalignment and poor model specification are more likely to negatively affect phylogenetic accuracy among fast-evolving sites than substitution saturation, corroborating previous theoretical findings based on less comprehensive datasets (6).

Results obtained using the Hug et al. dataset show that the phylogenetic signal present within slow-evolving sites is less consistent than that found within fast-evolving sites, especially for short-branched bipartitions. This observation is corroborated by analyses on simulated sequence alignment data, using a predetermined phylogeny and a known evolutionary model. This strongly suggests that the reported results are not artifacts due to unaccounted-for evolutionary processes, misalignment, or model biases. Among the simulated alignments, the fastest-evolving sites, expected to have experienced substitution saturation, still provided more accurate phylogenetic reconstructions than slow-evolving sites. The underperformance of highly conserved sites is accentuated in large scale phylogenies as bipartition branch lengths are skewed towards fewer substitutions per site (3, 8, 26). It follows that, as taxonomic sampling continues to increase, faster-evolving sites become more necessary for reliable phylogenetic reconstruction. These results challenge the assumption that the slower a site accumulates substitutions, provided it is not invariable, the better it is suited to reconstruct deep phylogenies (2, 27). Subjective alignment trimming strategies can generate phylogenies with significant differences in key bipartitions and can also impact the extant phylogenetic signal by affecting estimated model parameters (Table 1). The discussed results are not aimed towards proposing a new Tree of Life or evaluating any specific Tree of Life hypotheses, but rather, test the merits of current alignment trimming strategies and demonstrate how subjectively trimming fast-evolving sites may impair progress in these endeavors.

## Methods

### Hug et al. dataset

Hug et al. (2016) published a 3,083 taxa two-domain Tree of Life, proposing Eukarya as sister to Asgardarchaeota, and both within the TACK superphylum. The tree was reconstructed from a 2,596 sites super-matrix resulted from concatenating 16 ribosomal proteins, without trimming of fast-evolving sites or gene partitioning.

Aligned site positions were classified into twelve gamma-distributed substitution rate categories (α = 0.803) using IQTree’s parameter “-wsr” (10). Sites from each SRC were repeated *in tandem* to obtain the same length as the published alignment so that every RSAP has the same number of informative sites. RSAP phylogenies were reconstructed using IQTree with LG+G and pairwise distances between tree topologies were measured using Robinson-Foulds distances as implemented in IQTree.

During analysis of different alignment partitions to reconstruct phylogenies without slow and/or fast-evolving sites we considered sites from SRC1 to SRC4 as slow, thus removed from its respective analysis, while sites from SRC10 to SRC12 were considered as fast, and also removed from its respective analysis.

### Sequence simulation

Simulated sequence alignments containing 1,000 taxa were generated using Indelible software (28). A random tree topology was generated using the ETE Toolkit (29) and internal and terminal branch lengths were independently assigned from a gamma distributed branch length obtained from Hug et al. phylogeny. Shape parameters of gamma distributed internal and terminal branch lengths are 0.7581720 and 1.509421, respectively. Sequence simulations were performed with 100 replicates under an LG model with 12 gamma distributed site-specific substitution rate categories, no invariant sites, no indels, and same amino acid frequency as the Hug et al. dataset (control file available in Dataset S1).

### Phylogenetic analysis

All phylogenies were reconstructed using IQTree (12), and when necessary automatic model selection was performed using the “-m MFP” parameter. Ultrafast Bootstrap samples were obtained using the UFBoot method (11) and pairwise Robinson-Foulds distances (13) between trees were both estimated using IQTree.

RSAPs were generated by replicating sites from its respective SRC until the same length of the whole alignment is obtained. This was done to normalize the number of informative sites present in different SRC.

## Acknowledgements

This work was supported by the Simons Collaboration on Origins of Life grant #339603 to G.P.F.

## Supporting Information

**Figure S1.**
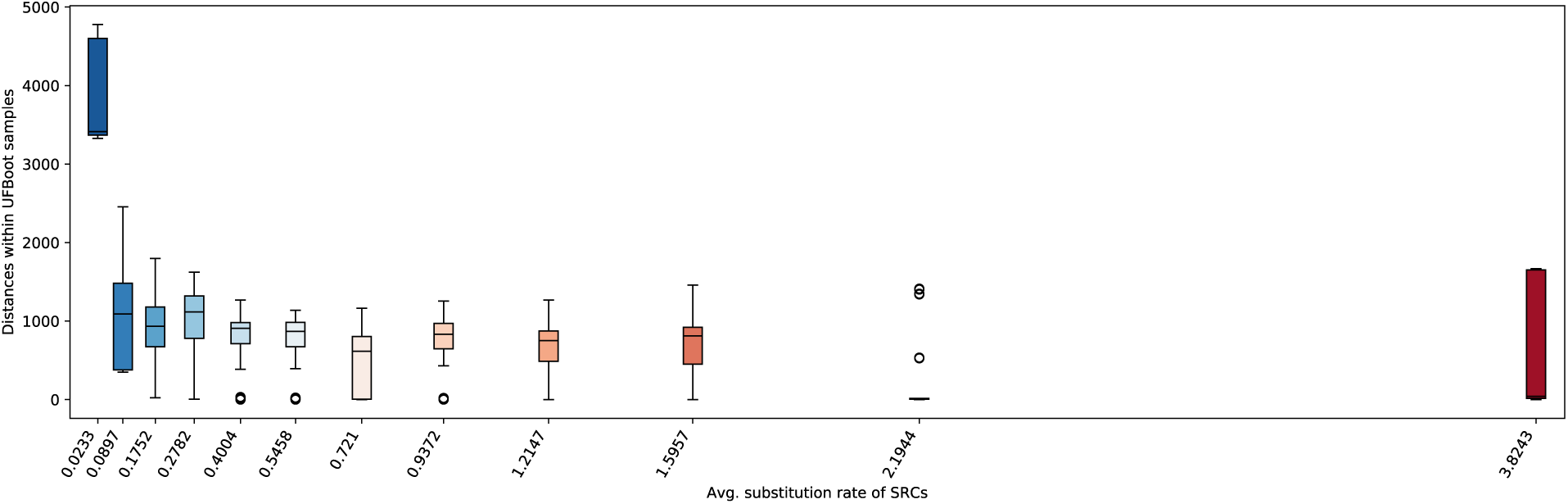
Boxplots representing pairwise RF distances among UFBoot samples within each RSAP from the Hug et al. alignment. Boxplot positions along the X axis represent the average site specific substitution rate of its sites.

**Figure S2.**
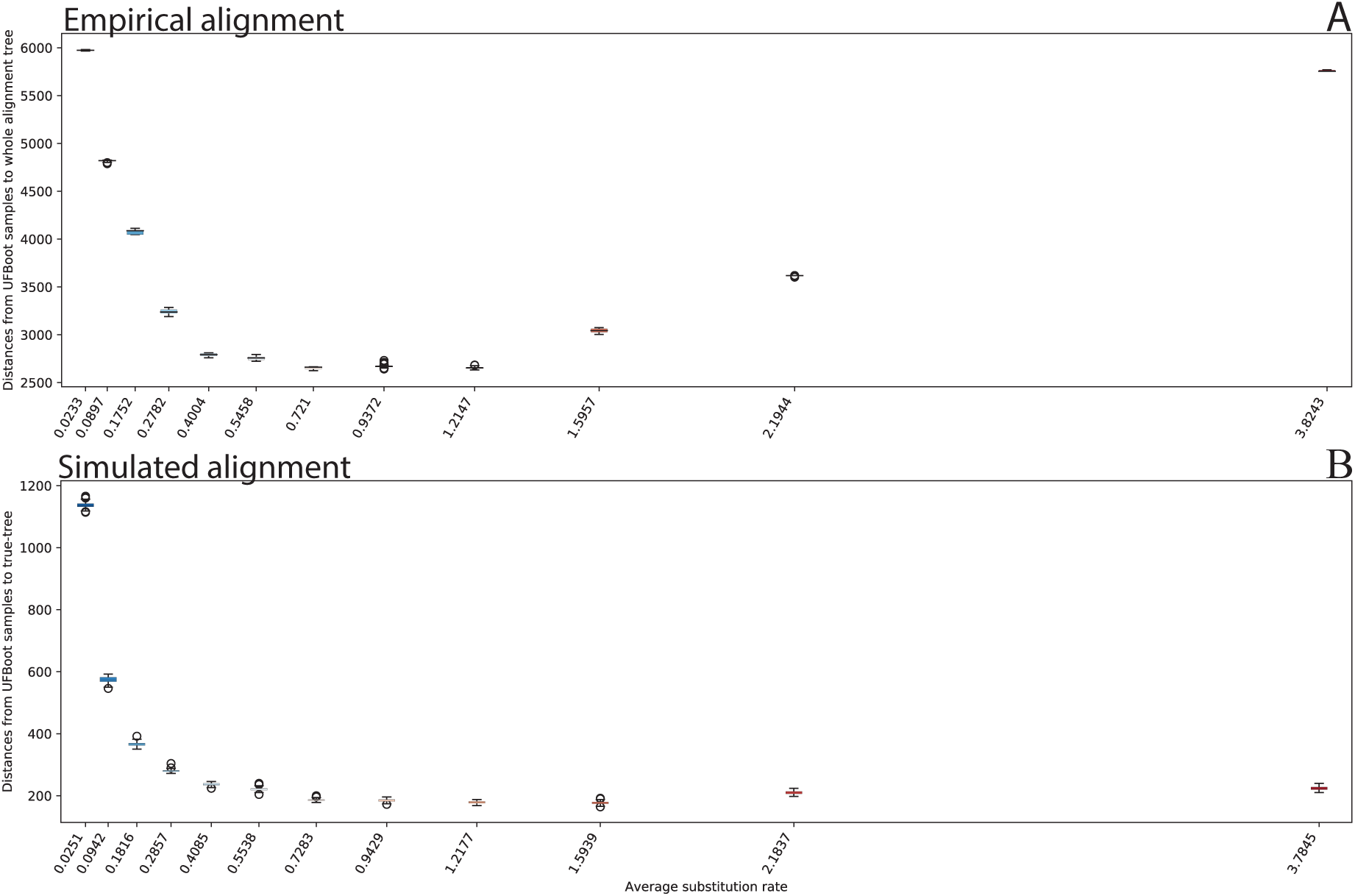
Boxplots representing pairwise RF distances between UFBoot samples from each RSAP versus its reference overall phylogeny. (A) RF distances between UFBoot samples and the Tree of Life topology reconstructed from the whole alignment published by Hug et al.. (B) RF distances between UFBoot samples against the true topology used to generate sequence simulations. Phylogenetic reconstructions of simulated dataset were performed using the same evolution model as its generation, LG+G. Boxplot positions along the X axis represent the average site specific substitution rate of its sites.

**Figure S3.**
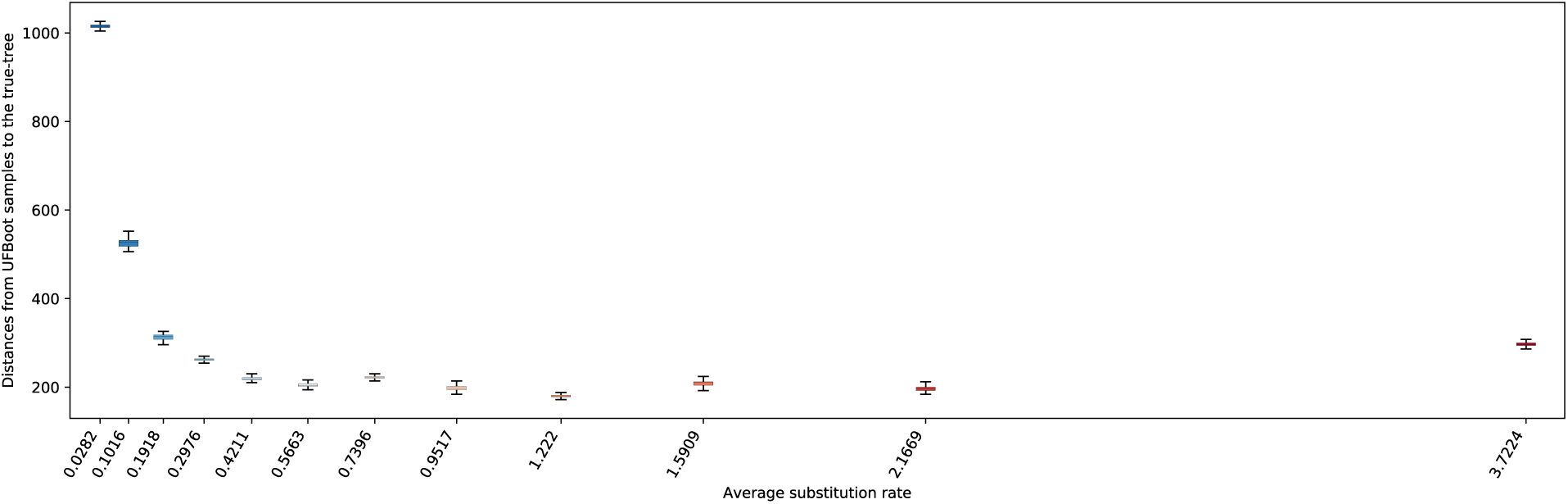
Boxplots representing pairwise RF distances between UFBoot samples of the simulation dataset reconstructed using Dayhoff+G model versus the true topology used to generate sequence simulations. Boxplot positions along the X axis represent the average site specific substitution rate of its sites.

**Figure S4.**
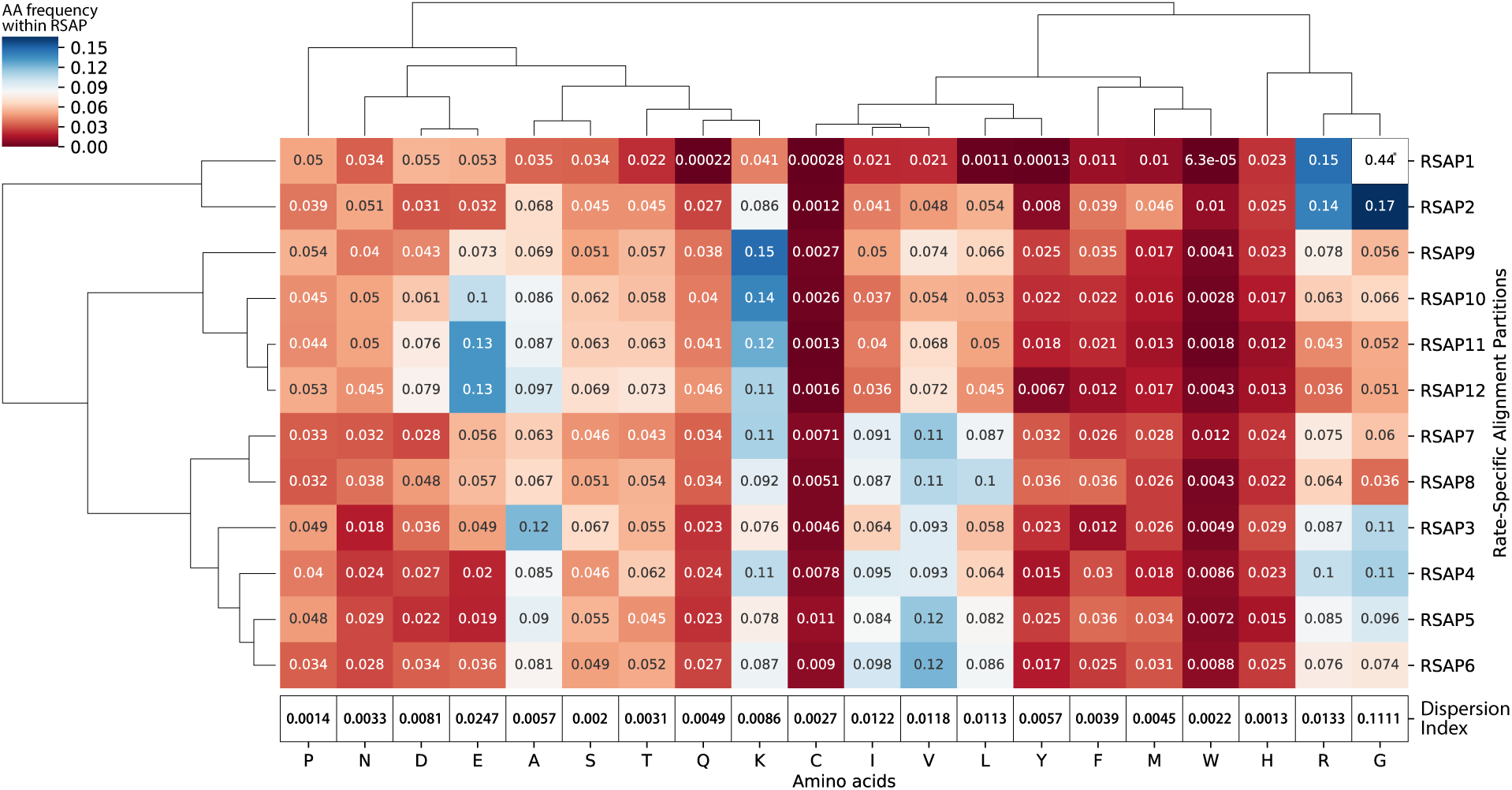
Hierarchical clustering of amino acids (X axis) and RSAPs (Y axis) using amino acid frequency correlations. The Dispersion Index row (defined as 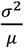) represents how much the frequency of each amino acid varies among RSAPs. Clustering was performed using complete linkage in both axes. *Glycine’s ratio within RSAP1 is out of scale to avoid color scale compression of the remaining frequencies.

**Figure S5.**
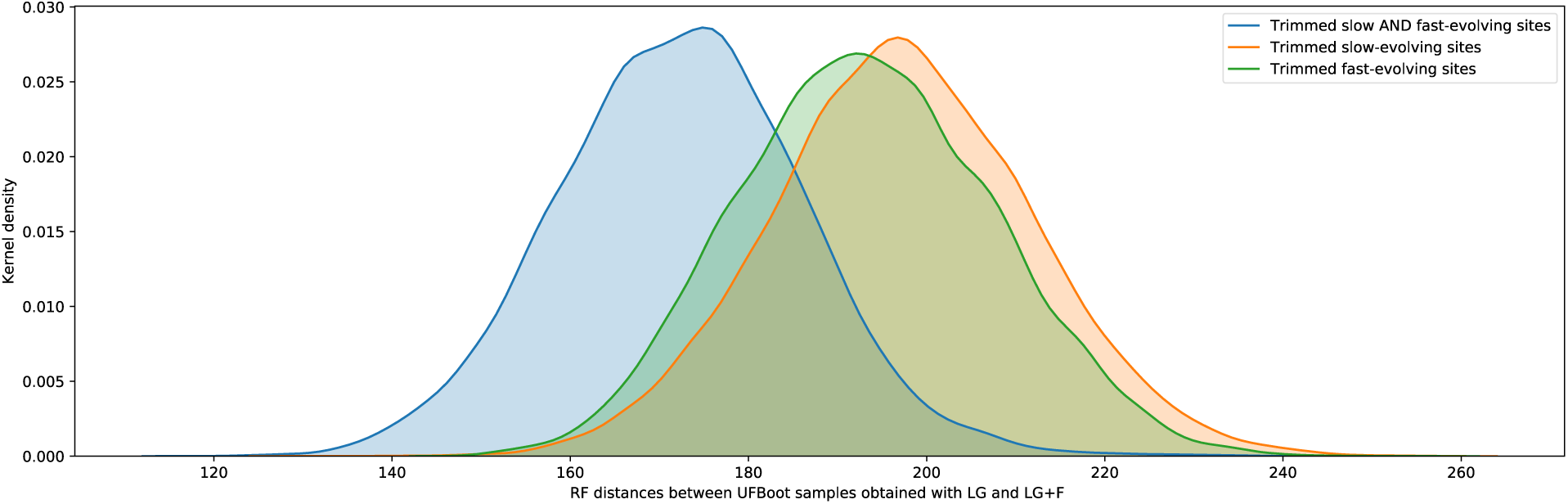
Distribution of RF distances between UFBoot samples obtained using two distinct evolutionary models: LG+G vs LG+F+G. In orange are RF distances between UFBoot samples obtained using the distinct evolutionary models with an alignment partition with trimmed slow-evolving sites (i.e., SRC1 to SRC4). Similarly, in green is the equivalent RF distances obtained with an alignment partition with trimmed fast-evolving sites (i.e., SRC10 to SRC12). Distances obtained using an alignment partition with trimmed both slow and fast-evolving sites are represented in blue.

**Figure S6.**
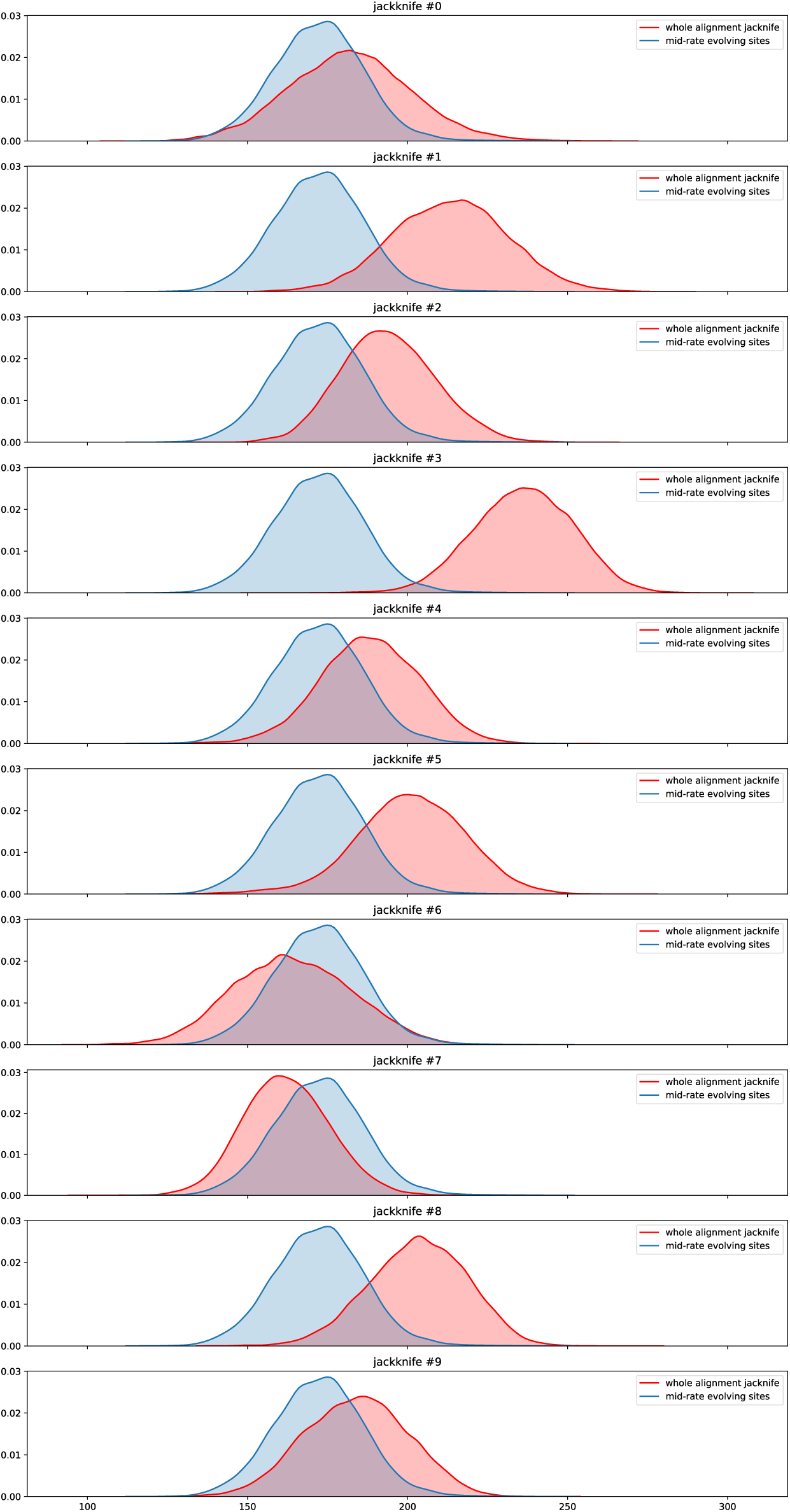
Distribution of RF distances between UFBoot samples obtained using two distinct evolutionary models: LG+G vs LG+F+G. Distances obtained using the whole alignment are represented in red, while distances from an alignment partition with trimmed both slow and fast-evolving sites are represented in blue. The whole alignment partition was jackknifed down to 1,420 sites, same length as the trimmed partition. Each subfigure represents a distinct random jackknife of the whole alignment.

